# On the status of Atlantic cod in the Oslofjord

**DOI:** 10.1101/2025.09.16.676469

**Authors:** Øystein Langangen, Stein Kaartvedt, Ketil Hylland, Joël M. Durant

## Abstract

There are serious concerns about the status of the Atlantic cod (*Gadus morhua*) in the Oslofjord region. However, quantitative estimates of the cod biomass are scarce. Typically, analysis of juvenile fish has been used to assess the state of the cod, and datasets that contain recruited individuals tend to be limited. We here quantify the biomass, with an estimate of uncertainty, over the last 25 years based on a large dataset of commercial landings. By accounting for the number of boat-landing events, we can account for the variation in effort over this period. Our analysis indicates that cod biomass has been reduced by more than 80% from a peak in 2004-2005 to 2018. After 2018, the biomass was further reduced below 10% of the previous levels, but these estimates are more uncertain. Given large public interest and a recent decision to close large areas in the Oslofjord for fishing, our results are highly topical and can serve as a baseline for further investigations into why the cod has collapsed in this area, as well as a contrast to potential future recovery.

## Introduction

The Atlantic cod (*Gadus morhua*) is an iconic fish species with immense historical, cultural, and socio-economic value (Kulatska et al., 2025; Kurlansky, 1997). The catches of cod in the Atlantic Ocean, including the North Sea region, have historically been high (Kulatska et al., 2025). However, the North Sea cod has recently likely undergone a regime shift, with the southern population being in a depleted state while the Viking population is recovering (Cecapolli et al., 2025). Similarly, the spawning stock of cod in the Kattegat is currently at very low levels, and recruitment is low (Kulatska et al., 2025). In Norway, cod management is delimited, depending on the location. Cod found within the 12 nautical miles of the coast are considered coastal cod and managed as such. The Oslofjord region (Fig.1) is located adjacent to the North Sea and linked to the Kattegat through, e.g., the Baltic Sea outflow into the Skagerrak (Gustafsson, 1997). In fjords along the Skagerrak, including the Oslofjord, there are at least two genetically distinct populations of cod: a *North Sea ecotype* typically distributed more offshore but also inside fjords and a *fjord ecotype* more tightly linked to the fjords (Barth et al., 2019; Knutsen et al., 2018). Cod in the Oslofjord is also potentially influenced by cod from the Kattegat, as tagging experiments have documented migration of adult cod from the Kattegat into Skagerrak (Righton et al., 2010). The cod in the Oslofjord, which seems to consist of more than 70% *North Sea ecotype*s for the fully recruited fish, has experienced a decline in catches over the last two decades (Moland and Strandli, 2021). This decline in catch is co-occurring with a generally reduced fishing effort in the region, making the link between catch and biomass of cod difficult to ascertain. In addition, the reduction in catches is only partially consistent with juvenile (0-group) data (Knutsen et al., 2022) showing relatively high abundances in some years. It is well-documented from other regions that mortality in cod is high and variable, also after the 0-group stage (Bogstad et al., 2016). It is hence important to obtain a better quantification of the changes in cod over time in the Oslofjord, based on recruited fish. Such quantification is topical due to large public interest in the recent negative trends for the Oslofjord cod, and a recent political decision to introduce full fishing ban zones in the region from 1^st^ of January 2026 (See Fig.1 and Press release in Norwegian^1^). In general, the Oslofjord has experienced a range of environmental problems, including eutrophication, reduced oxygen concentrations, coastal darkening, and increased sea temperature (Moland and Strandli, 2021; Ramon et al., 2025; Rogers et al., 2011). Climate warming is expected to impact cod recruitment negatively in this region as it close to the southern limit of the cod distribution(Drinkwater, 2005). Documentation of the quantitative changes that have occurred for cod is a first step to improving the status of the Oslofjord ecosystem. To this end, we use data from the Directorate of Fisheries based on reported landing tickets for the period 2000-2024 to quantify the catch, construct a proxy for effort to better link the catch data to biomass with uncertainty, and investigate how the size structure of the cod in the catch develops over time.

**Figure 1.**
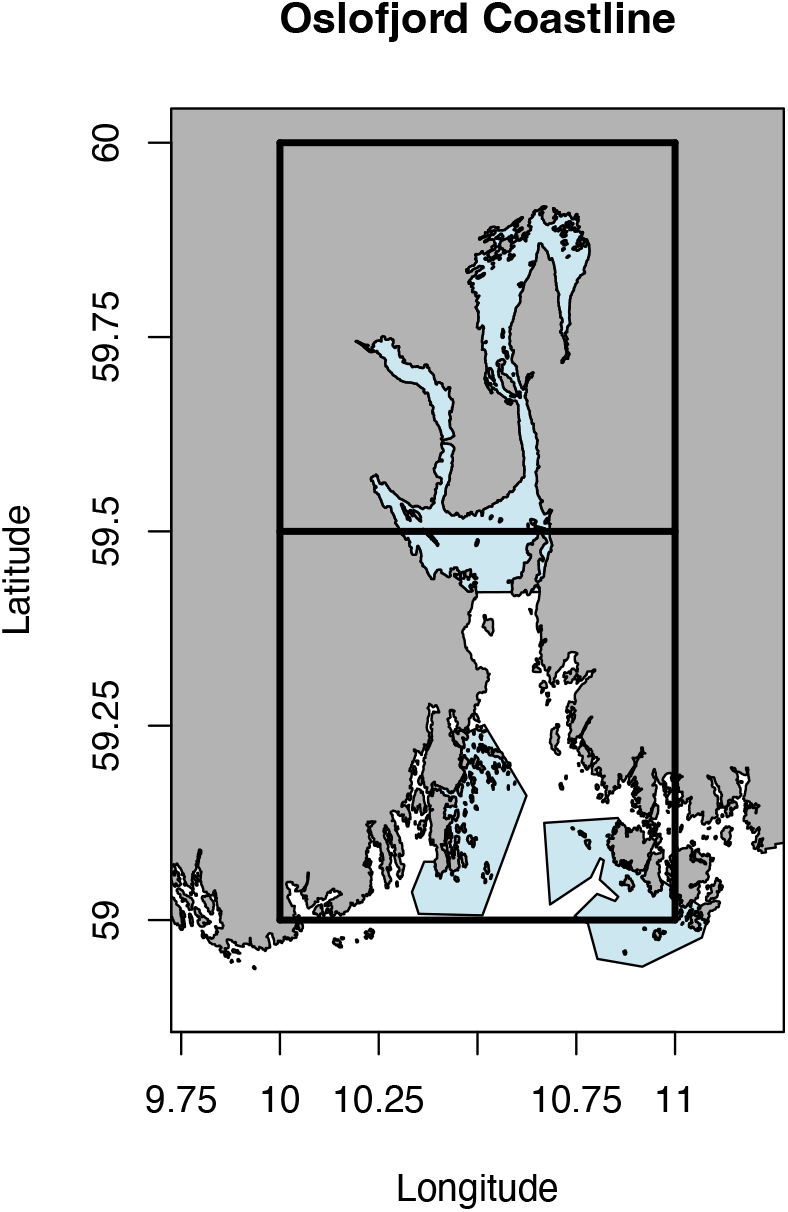
Overview of the Oslofjord region. The boxes show the extent of the statistical reporting areas, mainly covering the Oslofjord. Blue shaded polygons indicate proposed no-fishing zones redrawn from the recent press release (on page 3).

## Materials and Methods

### Data

Data for commercial landings is openly available from the Norwegian Directorate of Fisheries (www.fiskeridir.no) for the period 2000-2024. The landings are reported with main catch areas, with the statistical locations 9-20 and 9-22 representing the Oslofjord (Fig.1). The data for these two locations consist of almost 1.1 million entries. Northern shrimp (*Pandalus borealis*) is the most abundant species in the data, with more than 250.000 entries, followed by cod with more than 188.000 entries. For the cod data, the main gear reported was shrimp trawl (ca. 131.000) and set gillnets (ca. 36.000), i.e., parts of the reported landings are from bycatch in the shrimp fisheries. Catches of cod are reported over the whole year and are dominated by cod gutted without head (ca. 172.000), but also contain live cod (ca. 7800), gutted with head (ca. 4400) as well as roe (ca. 2300).

### Statistical analysis

We quantify the yearly catch in the region by summing up the landed weight (“Rundvekt”) for each landing ticket in the two statistical areas. To estimate the uncertainty, we performed a bootstrap by resampling boats through their vessel ID, within each year, with replacement. In 2000, the vessel ID were not available, precluding estimates of uncertainty.

The catch not associated with a vessel was removed from the input to the bootstrap and added to the bootstrap samples without error to avoid bias. The relationship between catch (C) and biomass (B) can be approximated by C = qEB, where q is the catchability and E is the effort. We here assume that q is constant over time (but see SI), and we used landing events, i.e., distinct vessel ID and landing date within each year, as a proxy for effort. Only vessels that reported cod at least once in the year were included. We thus implicitly assume that each fishing trip typically lasted only a single day; this is likely often the case for smaller boats (our results were robust to a test where only data from vessels smaller than 15 m were retained, results not shown).

By calculating the catch per landing event (i.e., catch per unit effort, CPUE), we construct a proxy for cod biomass. To estimate the catch and CPUE with uncertainty, we performed a bootstrap based on resampling the boats with replacement within each year and estimated the catch and effort for each sample. For this calculation, we did not include the catch and associated (but unknown) effort that were not linked to a vessel ID. Major management actions were introduced in 2015 (sorting grid for shrimp fisheries) and in 2019 (introduction of fishing ban zones), which likely affected the catchability of cod (q). To quantify these effects, we compared the CPUE between different gear types (shrimp trawl vs other) in different periods (see SI for details), and we calculated a corrected CPUE for the period after 2015. The estimated CPUE after 2019 is based on limited data and should be interpreted with care.

The data set moreover contains information on the size class of the catch. This size information is linked to the commercial value of the catch and is not always consistent across the years (Sletten Hopland and Aasheim, 2023). Nevertheless, size information is highly valuable as large individuals contribute disproportionately to, e.g., reproduction (Kopf et al., 2024), and size structure can therefore inform on the health of fish stocks. We analyzed the size classes reported for cod landed as gutted without head product class, which typically are reported in weight ranges <1 kg, 1-2 kg, 2-4kg, 4-7 kg, and > 7 kg. The latter class was only present in the data after 2004. As a proxy for the presence of large cod in the Oslofjord, we used the landed biomass of cod > 7 kg relative to total catch for the product class.

## Results

The calculated catch over time in Oslofjord is shown in Figure 2. The catch increased from around 150.000 kg in 2000 to a peak of around 250.000 kg in 2004 and then dropped steadily to about 84.000 kg in 2010, being fairly stable between 80.000-120.000 kg until 2015, with a further decline to low levels < 5000 kg in 2021 and 2024. This trend is somewhat different for the CPUE, with more stable CPUE largely between 20-30 kg cod per landing event for the years before 2015 (mean 24 kg), and a sharp decline to around 6 kg per landing event in 2018 and down to around 3 kg per landing after 2020. Analysis of the CPUE from different gear types in different seasons indicated that the catchability of shrimp trawls was reduced after 2015 compared to before. Correcting for this alteration in catchability increases the biomass estimates somewhat (Figure 2). We did not observe any clear trends in the relative presence of large cod in the landings (Figure 2).

**Figure 2.**
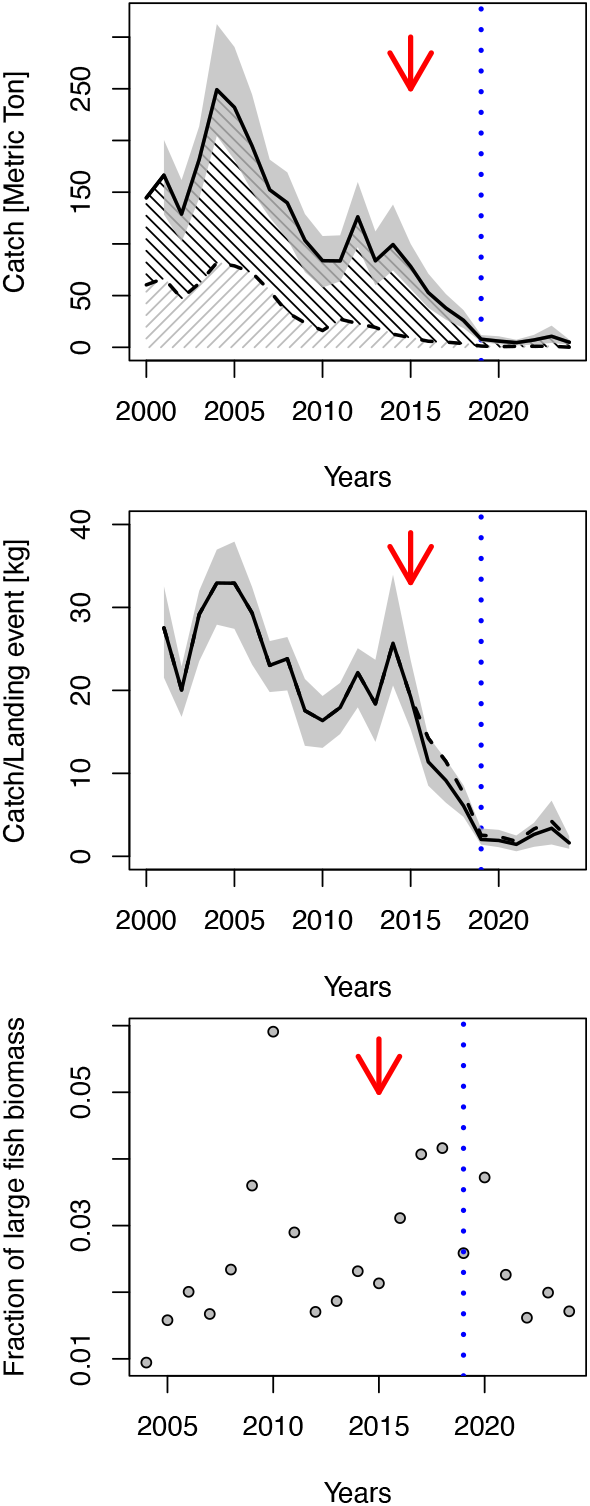
Total cod catches (upper panel, solid line) with a 95 % confidence interval (CI, grey shaded). The black hatched area indicates the landings from shrimp trawls, while the grey hatched area indicates landings from other gear types. The proxy for cod biomass is shown (central panel, solid line) with a 95% CI (Grey shaded), while biomass proxy corrected for altered catchability is shown (dashed line). The relative fraction of the biomass of large fish is shown in the lower panel. Red arrows indicate the introduction of separation grids for shrimp trawls, and blue dotted vertical lines indicate the introduction of fishing ban zones (2019). Results after 2019 must be interpreted with caution as there were limited landings in this period.

## Discussion

We have used a large commercial dataset to estimate how the biomass of cod in the Oslofjord region changes over time. With these data, we have documented a dramatic decrease in cod landings with part of the reduction being caused by reduced effort (Fig.2). Importantly, we have quantified a cod biomass proxy in the Oslofjord, with estimates of uncertainty. Our analysis documents a strong and significant decline in the Atlantic cod in the Oslofjord region, with the biomass dropping from a peak in 2004-2005 to below 20% of these peak values in 2018. The biomass further declined after 2019, but due to changes in the management regime in this year, the biomass proxy for the later years is presumably less reliable. Nevertheless, a striking feature of the time series of cod biomass is a period of relatively constant biomass followed by a steep decline after 2015. Note that the drop in CPUE is also observed when analyzing subsets of the data with different gear used (Figure S1). A scientific survey covering one transect conducted four times a year since 2011 shows a comparable trend in abundance indices, with a sharp decline after 2015 (Hylland, 2025). However, what is causing this decline in biomass is currently not clear, and it would be highly valuable to build models that can account for different mechanisms and drivers behind strong declines in population biomass, cf. (Olsen et al., 2024). Such potential drivers include fishing mortality and climate warming (Sguotti et al., 2019), contaminants (Ono et al., 2019), and ecosystem interactions (Olsen et al., 2024). Another potentially important mechanism is related to oxygen limitation and reduced growth, as documented for Baltic cod (Brander, 2022; Neuenfeldt et al., 2020). Successful models for addressing such questions, might (would?) require a better understanding of the population structuring, dynamics, and interactions of the sympatric populations in the region. One promising approach, which has been applied to European sprat (*Sprattus sprattus*) in the greater North Sea and beyond, utilizes population dynamical aspects to quantify the population structure (Lindegren et al., 2022). A similar approach, potentially combined with more classical genetic assessments (Knutsen et al., 2018), may be applied to cod in the region.

Also cod in nearby areas are in depleted states. This includes the Southern North Sea cod, which has experienced a collapse since 2015 (Cecapolli et al., 2025; ICES, 2024), mirroring the collapse observed in the Oslofjord region (Fig.2). The cod in both the eastern and western Skagerrak coast has seen a declining biomass over time (Stock et al., 2025). The connection to the other, adjacent populations of cod should be investigated further. We note that a small but significant fraction of the offspring from the Southern North sea and the Viking populations may drift into the Skagerrak and potentially into the Oslofjord (Romagnoni et al., 2020). Adult fish, especially from the Kattegat region, are also known to migrate into the Skagerrak (Righton et al., 2010). Surprisingly, we did not detect a reduction in large fish relative biomass in the landings over time (Fig.2). It may be due to targeted effort towards larger and more valuable fish, size selectivity in the bycatch from shrimptrawls, or it may reflect a less profound role of large individuals in the dynamics of cod in the region.

This study is associated with some limitations. For example, the data analyzed are based on commercial catches, which will reflect human behavior in addition to biological status. Moreover, there is some uncertainty in the reported main catch area, due to potential imprecise reporting and some limits on the spatial coverage of the reporting areas (Fig.1). The bootstrap may not fully capture the uncertainty associated with the estimates, as boat-level variation is not the only contributing factor. Effort and catchability may vary between boats, fishing trips, seasons, and years. However, we argue that the size of the dataset will likely reduce the impact of these limitations and that we here present an urgently needed quantification of the status of Atlantic cod in the Oslofjord. This information is highly valuable for scientists, decision makers, and the public alike.It may serve as a baseline for the evaluation of future development and potential recovery of cod in the area, also being of general interest in assessing the effects of the fishing ban on collapsed cod stocks.

## Supplementary Information

### Analysis with different gear types

To qua ntify the impact of alteration in gear use after 2015, we compared the CPUE from shrimp trawling (cpue_st, gear type numbers: 50, 51, 52, 55, 56, 57, and 83) and other gear (cpue_other, Fig. S1). For the period before 2015, the cpue_other were on average 1.21 times higher than the cpue_st. This is likely caused by diPerences in catchability. After 2015, but before 2019 (the year fishing ban zones were introduced), the ratio increased to 1.35 on average, with a max of 1.5. Assuming that the catchability in the shrimp fisheries was altered after 2015 compared to before, but that the catchability of the other gear was unaltered until 2019, would indicate that the catchability was reduced to about 89% (1.21/1.35) to pre 2015 values. Using the higher estimates (1.21/1.5) indicates a reduction to about 80 % of pre-2015 values for the catchability. To get an upper estimate of the biomass proxy for cod, we used this value to scale up the CPUE for *all* gear types in the landings after 2015 by dividing the CPUE by 0.80. The results are shown in Figure 2.

**Figure S1.**
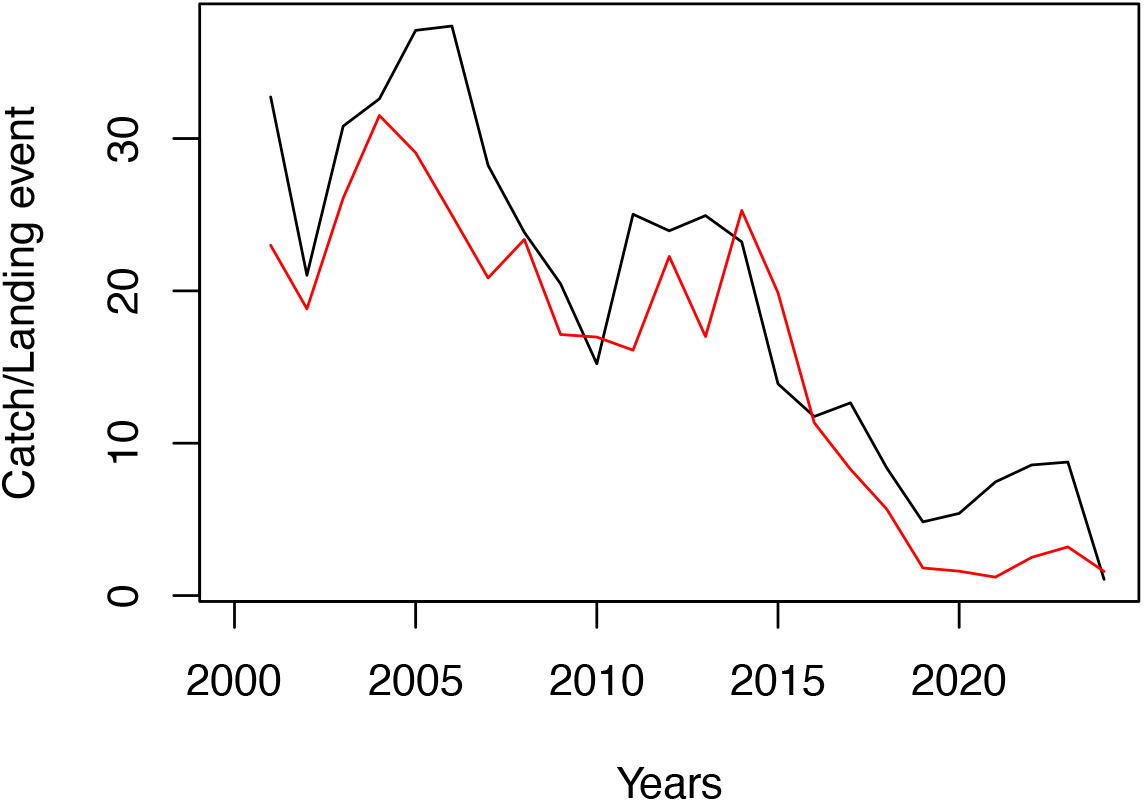
shows the CPUE based on data on shrimp trawls (red line) and CPUE from data with all other gear types (Black line).

